# Sensitive high-throughput assays for tumour burden reveal the response of a *Drosophila melanogaster* model of colorectal cancer to standard chemotherapies

**DOI:** 10.1101/2021.04.16.440211

**Authors:** Jamie Adams, Andreu Casali, Kyra Campbell

## Abstract

*Drosophila melanogaster (Drosophila)* models of cancer are emerging as powerful tools to investigate the basic mechanisms underlying tumour progression and identify novel therapeutics. Rapid and inexpensive, it is possible to carry out genetic and drug screens at a far larger scale than in vertebrate organisms. Such whole-organism-based drug screens permits assessment of drug absorption and toxicity, reducing the possibility of false positives. Activating mutations in the Wnt and Ras signalling pathways are common in many epithelial cancers, and when driven in the adult *Drosophila* midgut, it induces aggressive intestinal tumour-like outgrowths that recapitulate many aspects of human colorectal cancer (CRC). Here we have taken a *Drosophila* CRC model in which tumourous cells are marked with both GFP and luciferase reporter genes, and developed novel high-throughput assays for quantifying tumour burden. Leveraging these assays, we find that the *Drosophila* CRC model responds rapidly to treatment with standard CRC-drugs, opening the door to future rapid genetic and drug screens.

## Introduction

Colorectal cancer (CRC) is the third most common cancer and the second most frequent cause of cancer-related mortality^1^. With its high morbidity rate and incidence increasing year by year, CRC has become a global health problem. Chemotherapy is currently regarded as the standard treatment for patients with CRC, but it comes with many limitations, such as systemic toxicity, low selectivity and insufficient concentrations in tumour tissues^2^. Combined with the ability of tumours to develop resistance to currently used therapy regimes^3^, there is a need not only for alternative therapies that will increase chemotherapy treatment efficacy and reduce side effects, but also for new therapies that are less aggressive and more effective than conventional ones ^2,4^.

The process of identifying new CRC therapies requires the use of systems that are cost-effective, reliable and fast – preferably automated. Traditional approaches have employed high-throughput screening for small molecules that is based primarily in 2D or 3D cell culture, enzymatic assays or receptor binding assays. Such studies in 2D have relied on the use of CRC cell lines such as Colo 205, SW480 or CR4, and assaying for effects on key markers and genes required for growth, migration and invasion using a combination of immunostaining, basic cell morphology, viability and proliferation assays^5-8^. 3D studies in spheroids or organoids have provided the opportunity for more in-depth study of the effects of treatments on physical tumour characteristics, with less reliance on biological markers such as E-cadherin or Vimentin^9,10^. Increasingly complex manipulations of 3D *in vitro* techniques have allowed the control of a variety of biochemical and biophysical factors, while providing the capacity for real-time imaging with standard microscopy. While 2D and 3D cell-culture systems have evolved to enable more sophisticated high-throughput therapeutic screenings, the ease of use and fast-turnaround times of such *in vitro* studies comes at the cost of removing the rich biological environment found *in vivo*, alongside a lack of targeted therapy delivery and limited physical outputs. Consequently, the majority of positive hits identified through these types of *in vitro* screens, are later found to be ineffective and/or toxic in subsequent validation experiments in whole-animal models^11^.

*Drosophila melanogaster* are emerging as a powerful alternative platform for high-throughput drug screens. With a rapid life cycle and low maintenance costs, such whole-organism-based drug screening permits the assessment of drug absorption, distribution, metabolic stability and toxicity, reducing the possibilities of false positives. With nearly 75% of human disease-causing genes believed to have a functional homologue in the fly^12^, *Drosophila* are increasingly used as screening platforms for drug discovery and translational research^13,14^. There is a striking similarity between mammalian and *Drosophila* intestinal tracts, which makes them particularly suited for modelling CRC^15^. While on a gross anatomical level, the intestinal architecture differs between mammals and *Drosophila*, they are composed of similar cell populations. The intestines in each are maintained by proliferative cells that give rise to two types of post-mitotic cell types; absorptive cells called enterocytes with a characteristic brush border and secretory cells. Therefore, the *Drosophila* intestinal tract presents a ‘stripped down’ version of mammalian intestines that retains essential features of the stem cell biology and niche, which are among the essential features when modelling CRC.

A genetic powerhouse, the fly’s experimental tractability has identified a number of cancer initiating mutations and has led to the generation of a large number of *Drosophila* models for different cancers^16,17^. Combined mutations in the WNT and EGFR pathways are the most frequent initiating events of CRC^18^, and we and others have previously shown that it is possible to model CRC in *Drosophila* by inducing clonal activation of the Wnt and Ras signalling pathways in the adult midgut^19,20^. Intestinal epithelial ApcRas clones mutant for the negative regulators of the Wnt pathway, *Apc* and *Apc2*, and overexpressing the oncogenic form of Ras, UAS-Ras^v12^, form tumour-like overgrowths. We showed that these overgrowths reproduce many hallmarks of human CRC, including increased proliferation, a block in cell polarity and differentiation, as well as disrupted organ architecture. Furthermore, metabolic behavioural assays showed that these flies suffer a progressive deterioration in intestinal homeostasis^19^.

The behaviour of mutant clones generated in the adult *Drosophila* midgut is commonly analysed by dissecting and imaging midguts, and quantifying parameters such as the number of clones, the number of cells in a clone and the clonal area manually using standard imaging analysis software^19,21,22^. As these methods are laborious, they cannot be done on a rapid, large scale. Here, we have developed two approaches to make the analysis of tumour burden in the *Drosophila* CRC model high-throughput, which could be easily adapted to other cancer models. First, we have developed custom built macros to be used in conjunction with Fiji, a widely available distribution of ImageJ for Life Science, which allow for the automated analysis of the total area of the midgut taken up with clones, as well as the average clone size. We demonstrate that clone size is the most robust determinant of whether a clone is behaving as tumour or not. Secondly, having introduced a second reporter into the system, with clones now labelled with both GFP and luciferase, we show that the overall tumour burden can be rapidly and cheaply assessed by assaying whole fly lysates for luciferase. Finally, we show that after feeding drugs to the flies in their food, these assays can be used to assess tumour burden accurately and rapidly, and demonstrate a robust response of the *Drosophila* CRC model to conventionally used CRC therapies. This study acts as a proof of principle to show that this model has the capability to be used in a high-throughput manner for future small compounds screens, and generates quantitative measurements of tumour characteristics that allow more in-depth characterisation of the efficacies of compounds.

## Methods

### Clone generation

MARCM clones were generated by a 1-h heat shock at 37 °C of 2–5-day-old females and were marked by the progenitor cell marker escargot (esg) Gal4 line driving the expression of UAS GFP.

### Genotypes

*yw hsp70-flp; esg-Gal4, UAS-GFP, UAS-Ras*^*V12*^*/CyO; UAS-Luciferase FRT82B, Gal80/TM6b* flies were crossed with *yw hsp70-flp; Sp/CyO; FRT82B, Apc2*^*N175K*^*Apc*^*Q8*^*/TM6b* flies to generate ApcRas clones. *yw hsp70-flp; esg-Gal4, UAS-GFP/CyO; UAS-Luciferase FRT82B Gal80/TM6b* flies were crossed with *yw hsp70-flp; UAS-Sna/CyO; FRT82B/TM6b* flies to generate Snail clones and *to yw hsp70-flp; If/CyO; FRT82B/TM6b* flies to generate GFP clones. Apc2^N175K^ is a loss-of-function allele, Apc^Q8^ is a null allele, UAS-Ras^V12^ is a gain-of-function transgene, and UAS-Sna is a wildtype transgene. All balancers (Cyo and TM6b) were compound balancers. UAS-Sna was a gift from Justin Kumar. All other stocks were obtained from Bloomington Stock Centre.

### Drug administration

5-Fluoro-5’-Deoxyuridine (Sigma Aldrich, F8791) and Oxaliplatin (Sigma Aldrich, O9512) were dissolved in DMSO and added to molten fly food – ensuring temperature of molten fly flood did not exceed the thermostabilities of compounds. DMSO alone was added to molten fly food as a control. Drugs were diluted to a concentration of 100 µM, as has been shown to be effective in other studies where compounds are fed to *Drosophila* cancer models^23^.

### Staining and antibodies

Adult female flies were dissected in phosphate-buffered saline (PBS). All whole digestive tract was removed and fixed in PBS and 4% electron microscopic-grade paraformaldehyde (Polysciences, USA) for 35 min. Samples were rinsed 3 times with PBS, 4% bovine serum albumin (BSA), and 0.1% Triton X-100 (PBT-BSA) and incubated with the primary antibody overnight at 4 °C and with the secondary antibody for 2 h at room temperature. Finally, the samples were rinsed 3 times with PBT-BSA and mounted in DAPI-containing media (Vectashield, USA). All the steps were performed without mechanical agitation. Primary antibodies were goat anti-GFP (1:500; Abcam, ab6673) and rabbit anti-Laminin (1:500, Abcam, ab47651). Secondary antibodies were from Invitrogen (USA). Phalloidin (Sigma, USA, P1951) was used at 5 µg/ml. Confocal image were acquired with a Zeiss LSM 880. Images were analysed with the Fiji software [National Institutes of Health (NIH) Bethesda, MD] and assembled into figures using Fiji, the Adobe Photoshop software, and Microsoft Powerpoint.

### Luciferase assays

Luciferase assays were performed using the Dual-Luciferase(R) Reporter Assay System. Flies were squashed using a 200µl pipette tip into a Passive Lysis buffer and samples loaded into 96-well plates. Luciferin substrate (LARII) added and bioluminescence read on a Varioskan plate reader.

### Image analysis

The macros presented in this study are for use with ImageJ (https://imagej.net/Fiji/Downloads). ImageJ is an open source imaging processing and analysis platform, originally developed at the National Institutes of Health (Bethesda, Maryland, USA). We recommend using ImageJ Fiji distribution. In principle Fiji should run on all major computer platforms, including Microsoft Windows, Mac OS and Linus.

The macros are briefly described below and can be found here: https://github.com/Jadams94/AutomatedGutQuant

BatchGutGFP%CoverageMacro: This macro measures the overall area of GFP positive pixels and the overall area of an object in the red channel of an RGB image. These area measurements can then be used to calculate the overall GFP% coverage of an object within an image.

BatchGutGFP%CoverageMacro: This macro measures the overall area of GFP positive pixels and the overall area of an object in the red channel of consecutive RGB images within a specified folder. These area measurements can then be used to calculate the overall GFP% coverage of an object within each image.

GFPMaskAreaQuantMacro: This macro measures the area of each individual GFP positive mask of an RGB image.

These macros should be downloaded onto your computer and dragged and dropped to the Fiji bar to be used. The purpose and directions for use of each macro are incorporated within each file.

### Statistical analysis

All statistical analyses and graphical representation were carried out using GraphPad Prism 8.

## Results

We have previously shown that, by means of the MARCM technique^24^, we can induce GFP-marked Apc-Ras clones in the intestinal stem cells of adult midguts. After 4 weeks, these clones form tumour-like overgrowths that share many similarities with human CRC (Fig.1a and ^19^), generating an excellent model in adult *Drosophila* for CRC^19^. In order to be able to systematically quantify tumour growth, we recently generated a new CRC model that expresses a UAS-luciferase transgene^25^, in addition to UAS-GFP, in the ApcRas background^26^. This model allows us to not only visualise cells using GFP, but also to exploit the highly sensitive luciferase-based assay to test for tumour burden, a process which is simple, quick to perform and provides rapid results. First, whole flies are homogenized directly into a lysis buffer, then the luciferin substrate is added, and finally the amount of luciferase activity is read using a luminometer (Fig. 1b)^27^. As luciferase readings can be performed using a microplate reader capable of measuring renilla luminescence, this process can be scaled up considerably, facilitating the analysis of many conditions in parallel.

**Figure 1.**
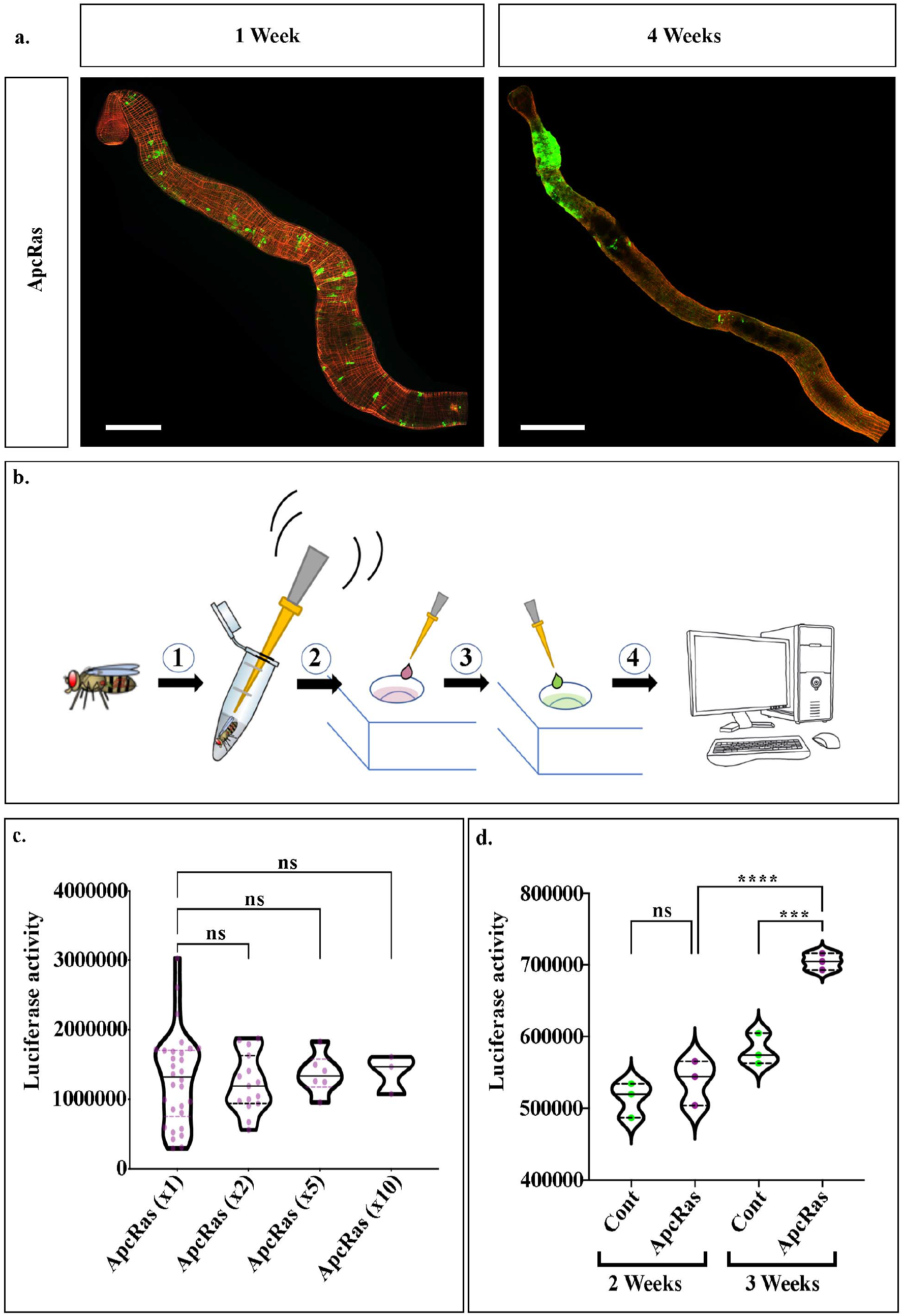
Assessing tumour burden by assaying for luciferase activity in whole flies. (**a**) Adult midguts dissected either 1 week (left), or 4 weeks after clone induction (right), stained for GFP (green) and phalloidin (red). By 4 weeks ApcRas clones form large tumours in the anterior region of the midgut. (**b**) Schematic showing how luciferase assays are performed. Flies are homogenised in a passive lysis buffer, which is then pipetted into a 96-well plate. Luciferin is added and the resultant light is registered and processed via a plate-reader (**c**) Luciferase assays performed on 30 flies individually, in pairs, as groups of 5 and in groups of 10, respectively. (**d**) Luciferase assays performed on 3 batches of 10 control flies with wildtype clones, or of flies bearing ApcRas clones. Graphs show violin plots with the median (solid line) and upper and lower quartiles (dashed lines). Statistical analysis (c) is one-way ANOVA and (d) a Kruskall-Wallis test **** P ≤ 0.0001; *** P ≤ 0.001; ns = non-significant. Scale bars = 500 μm.

As a first step towards setting up the conditions for screening whole flies for tumour burden using luciferase, we first separately assayed the luciferase activity in homogenates from 30 individual flies bearing ApcRas clones (Fig 1b). In agreement with the high variability we described in this CRC model^19^, we observed a high variability in the luciferase readings between each fly, and thus used these measurements to determine the average luciferase activity in a fly with ApcRas clones (Fig 1c). We next assayed the luciferase activity in homogenates taken from pairs of flies, groups of 5 flies, or groups of 10 (Fig 1c). We found that increasing the number of flies to be processed together, not only reduces the variability in the luciferase readings, but leads to an almost identical average readout for luciferase activity. Taken together, this suggests that a rapid assay where 3 groups of 10 homogenised flies are put together, should give as reliable results as 30 individual assays. Using these conditions, we compared control flies bearing wildtype clones with flies bearing ApcRas clones, and found that while they gave comparable luciferase readings at 2 weeks, at 3 weeks their luciferase readings were significantly higher, correlating with our previous observations of the timing of tumour development in this model (Fig 1d compare with ^19^).

We next sought to characterise in more detail how the whole fly luciferase readings relate to the behaviour of GFP positive clones of cells within the gut. As well as providing a more in-depth analysis of cell behaviour in different conditions, we reasoned that this could be used as a secondary screen to validate hits from a luciferase screen. This process, which we established when we developed and characterised the ApcRas model^19^, involves dissecting guts from mated females at 2 and 3 weeks after clones have been induced, fixing and staining for GFP to visualise the clones, and phalloidin, which labels F-Actin and shows the morphology of the midgut. As shown previously, control clones labelled just with GFP and luciferase, form small clusters dispersed throughout the gut, that don’t appear to increase much in size (Fig. 2a). In contrast, ApcRas clones disappear over time, with the few that survive forming tumour-like overgrowths (Fig. 1a, 2a and^19^). Until recently, we quantified tumour burden by imaging each midgut, stitching the images together manually (midguts can be up to 6mm long^28,29^), and then used standard functions on Fiji to outline and calculate the area of each clone. While this method provides highly accurate parameters to describe the behaviour of clones, it is extremely laborious and can also introduce user bias.

**Figure 2.**
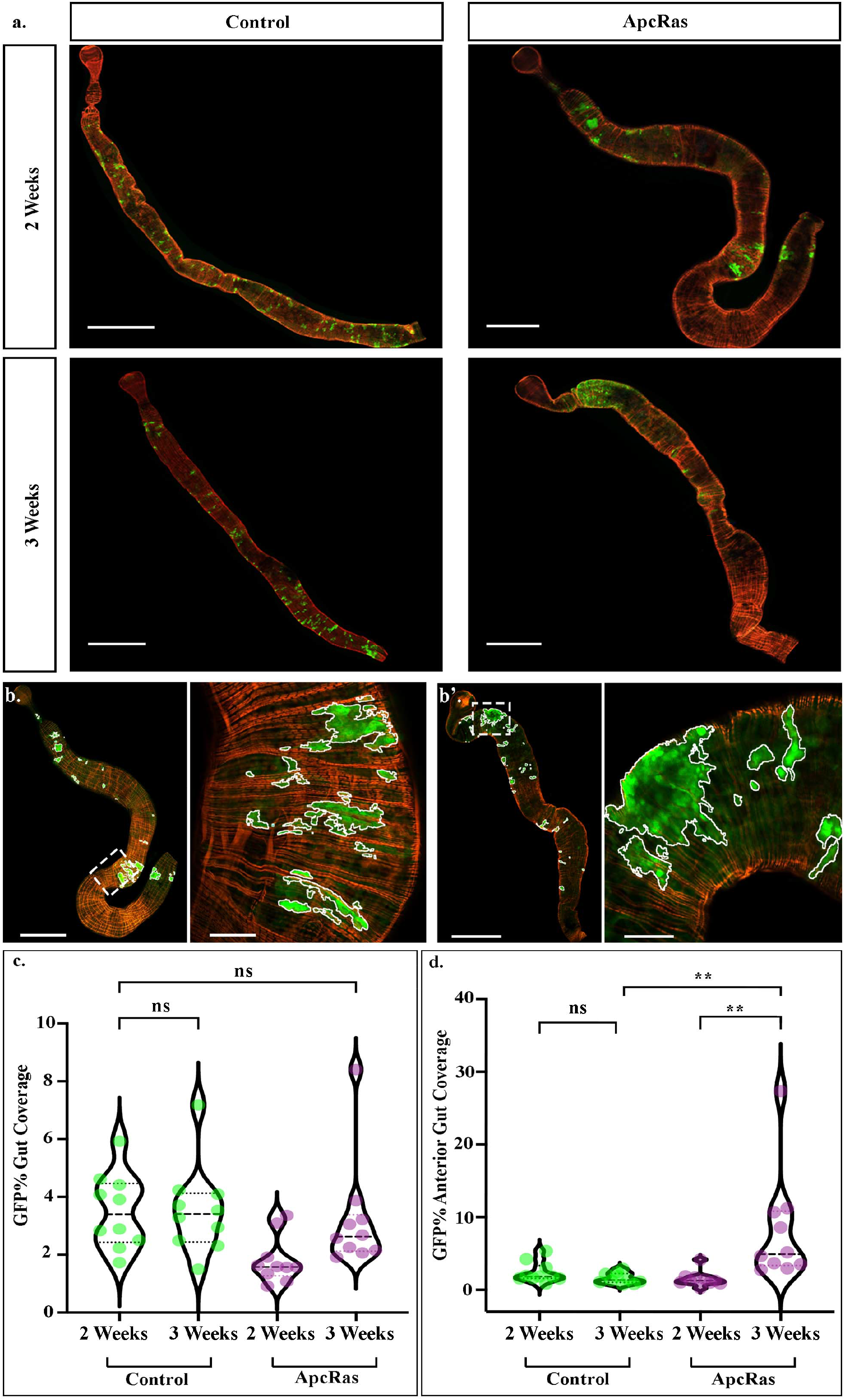
Tumour burden can be analysed automatedly across the entire midgut. **a, b** dissected adult midguts are stained for GFP (green) and phalloidin (red) (**a**) GFP labelled clones 2 and 3 weeks after clone induction in Control and ApcRas conditions. (**b**) GFP positive areas across whole guts can be accurately selected using custom automated scripts. Blown-up image sections are outlined by dashed white-lines (**c**,**d**) Quantifications of the GFP% coverage of the anterior-portion of the midgut (**c**) and GFP% coverage of the entire midgut (**d**) of control, and ApcRas clones at 2 and 3 weeks, n = 10. Graphs show violin plots with the median (solid line) and upper and lower quartiles (dashed lines). Statistical analysis is Kruskall-Wallis test. ** P ≤ 0.01; ns = non-significant. Scale bars = 500 μm (whole guts) and 50 μm (close-ups in b).

Moving towards a more rapid system for clonal analysis across entire midguts, we started to acquire tile-scans using a laser scanning microscope and image-stitching software, which enables the automated acquisition of a single continuous image of the entire organ. We have developed new tools for the analysis and extraction of reliable quantitative data from these tile scans. We have made these custom scripts publically available (see Methods section for details on how to download and use them). They are for use with ImageJ, an open source imaging processing and analysis platform, and allow the automated and accurate identification of GFP-positive areas along the entire midgut in seconds, as well as the quantification of various parameters such as area of each individual clone, or sum of the areas of all clones within a midgut (Fig. 2b-d). They also provide an option for a variable threshold which enables images of a range of qualities and microscopes to be analysed in a similar manner, allowing comparisons to be made across a variety of equipment.

As ApcRas clones are confined to the anterior portion of the midgut by 3 weeks after clone induction, we previously quantified tumour burden by calculating the % of the anterior portion of the midgut taken up by GFP positive clones^19^. We therefore tested our custom macros by measuring the total clone area of wild type or ApcRas clones within the anterior portion of the midgut, along with the area of the anterior portion of the midgut itself, enabling the percentage area of the anterior midgut covered by clones to be calculated automatedly. While this requires manual selection of the anterior portion of the midgut, the use of thresholding and the automated quantification of GFP positive area removes potential user bias and, with proper thresholding, selects GFP positive clones only in the current Z-plane.

While the percentage coverage of the anterior portion of the midgut presents a reliable metric for analysing changes to the tumour burden in the ApcRas model, this may not extend to other models where clones are more evenly distributed throughout the midgut. We therefore also developed a macro to measure the entire midgut area, enabling the calculation of the percentage of the entire midgut covered by clones. In order to further increase the high throughput capabilities of this model of CRC, we also developed a macro that quantified the GFP positive area, along with the whole area of objects in the red channel, of consecutive images. This allows extremely large batches of dissected guts, or even whole experimental conditions, to be processed in minutes. Interestingly, while the calculation of the GFP positive area in the anterior region of midguts is significantly higher in the ApcRas conditions than in controls (Fig. 2d), the percent of the entire midgut covered by GFP positive clones shows no significant difference (Fig. 2c). This suggests that while the percentage coverage of the anterior portion of the midgut may be a good metric for analysing the tumour burden in the ApcRas model versus wildtype, the percentage coverage of the whole midgut is not. Therefore, for a high-throughput screen for drugs altering the tumour burden in the ApcRas model, which may affect the distribution of clones across the midgut, or for measurements in any condition where clones are not restricted to one portion of the midgut, other parameters may be more informative.

This raises the question as to what quantifiable parameters may determine the difference between a wildtype control clone, a clone that has over-proliferated but not formed a tumour, and a tumour-like clone where cells have lost polarity and are undifferentiated. In previous work, we have shown that the overexpression of Snail in the intestinal stem cells leads to the over-proliferation of cells, but clones do not form tumours^26^. We therefore decided to look in detail at the cell morphology, and distribution of either control clones, clones over expressing Snail or ApcRas clones (Fig 3a-f), to identify quantifiable parameters that could be leveraged to distinguish what kind of clone has formed. We found that the tissue architecture, cell organisation and nuclei size are distinct between control clones (Fig. 3a,b), Snail clones (Fig. 3c,d) and ApcRas clones (Fig. 3e,f). The adult *Drosophila* intestinal tract shows high levels of cellular heterogeneity, as the epithelium is composed of a range of cell types, as well as cells of varying ages, as the epithelium undergoes continual cell turnover. Regardless of this high heterogeneity, the cell morphology and general tissue architecture in control clones and Snail clones is relatively consistent and well-organised (Fig 3a-d), while ApcRas clones exhibit poor cell organisation, large variation in cell morphology and shape, increased nuclei density within clones and a huge range of nuclei sizes – characteristics displayed in mammalian epithelial tumours (Fig 3e,f). A key difference between clones of cells which have overproliferated, versus tumour-like clones is an increase in the maximal nuclear size; the maximal nuclear size in control and Snail clones is 420 µm^2^, whereas nuclei in ApcRas are on average bigger and can be as large as 980 µm^2^ in diameter. While nuclei in ApcRas clones are more packed than in controls (Fig 3 compare e with a), this is also seen in over proliferating conditions (Fig. 3c). In contrast, while control and Snail clones are always found within a single layer (Fig. 3b,d), ApcRas clones are multi-layered and protrude into the lumen of the midgut (Fig. 3f). However, assessing whether clones have become tumour-like by their maximum nuclear diameter and extent of which they have become multi-layered or protrude into the lumen is very time consuming, as the midgut is a very thick tissue and it requires taking many z-slices in high resolution. Combining this with the need for tile scans to capture the entire length of the midgut, this suggests that while these parameters can be used to assess a few samples, it would not be practical to use them as outputs for a high-throughput screen.

**Figure 3.**
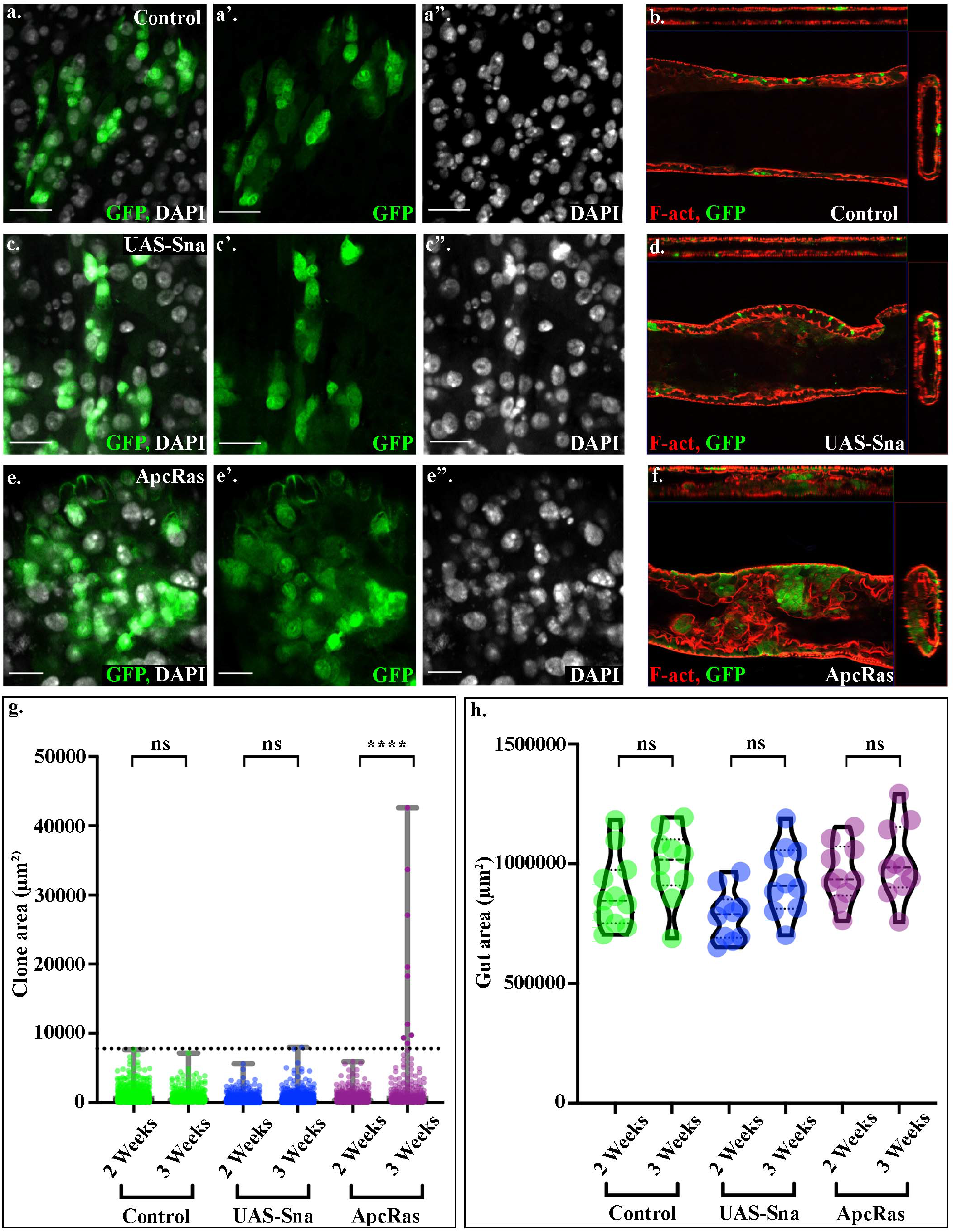
Distinguishing between control, over-proliferating and tumour-like clones. **a-f** representative clones from Control, UAS-Sna and ApcRas midguts stained for GFP (green) and Dapi **(a**,**c**,**e** white) or phalloidin **(b**,**d**,**f** red**). a**,**c**,**e** longitudinal sections and **b**,**d**,**f** orthogonal views of control (**a**,**b**) UAS-Sna (**c**,**d**) and ApcRas (**e**,**f**) midguts. While the lumen of midguts containing either Control or over-proliferating clones consist of a single-layered epithelium and are continuous and clear (**b**,**d**), there is multi-layering in the epithelium of midguts with ApcRas clones, which often have luminal-blockages and a discontinued lumen (**f**). **g**,**h** Graphs showing scatter plots with min and max (**g**, solid horizontal lines) and violin plots with the median (**h**, solid line) and upper and lower quartiles (**h**, dashed lines). (**g**) Quantifications of the area of clones found in midguts with either Control, UAS-Sna or ApcRas clones. Clones of over 7,800 μm^2^ (dashed line delineates this threshold) are only found in the ApcRas model. Statistics were performed on the number of clones found over the threshold. (**h**) The average area of the whole midgut is not significantly altered in any of the conditions. Statistical analysis is Kruskall-Wallis test. **** P ≤ 0.0001; ns = non-significant. Scale bars = 20 μm.

The formation of these large multi-layered tumours, suggested that tumour-like clones could be distinguished from controls and over-proliferating cells by size. Therefore, automated scripts were developed to measure clone area of all clones along the entire midgut. These showed that the large clones over 7,800 μm2, likely formed by the merging of multiple clones, are only seen in ApcRas conditions, and never seen in control or in Snail clones (Fig. 3g). This occurs without any significant changes in the overall gut area (Fig. 3h). Importantly, comparable results were seen whether these were analysed from a tile scan across 3 z planes of the midgut, or across a z-stack through the entire midgut. Taken together, these results suggest that the clone area is a robust and reliable readout for the presence of tumour-like clones in the midgut and can be assayed and compared using just 3 z planes.

Once the conditions for screening ApcRas flies for tumour burden had been established, we next wanted to test whether the *Drosophila* CRC model responds to drugs currently used to treat human tumours. We decided to test two drugs: Oxaliplatin and 5-Fluoro-5’-Deoxyuridine (5-Fluoro), two chemotherapy drugs that are standard treatments for patients with CRC^30^. Oxaliplatin is a platinum-based DNA crosslinking agent^31^, whereas 5-Fluoro is a Fluorouracil derivative which acts as an antimetabolite by blocking synthesis of thymidine^32^, and both act to block cell division. We reasoned that their efficacy in treating human CRC suggested that they would be good candidates to test our model.

Flies bearing ApcRas clones were induced in adult midguts and allowed to develop for 1 week. The flies were then transferred to new vials containing either food alone (Untreated), food mixed with Oxaliplatin or 5-Fluoro or with the additional control of Dimethyl sulfoxide (DMSO) alone, as this was used to dilute the drugs before adding to the food. The drugs were diluted to a final concentration of 100 µM within the food. Flies were left on the food with drugs or control food for 2 weeks, and then assayed for tumour burden by luciferase assays on whole flies (Fig. 4a). These revealed that treatment with either Oxaliplatin or 5-Fluoro leads to a large reduction in the overall tumour burden when compared to untreated flies, or flies fed with DMSO alone (Fig. 4b). This demonstrates that *Drosophila* CRC flies show a rapid and robust response to treatment with standard human CRC therapeutics. Moreover, this suggests firstly that the *Drosophila* CRC model may be a viable screening platform for drugs for treating human CRC. Secondly, as luciferase assays can be performed on whole fly homogenates in 96-well plates, this approach is scalable to high-throughput.

**Figure 4.**
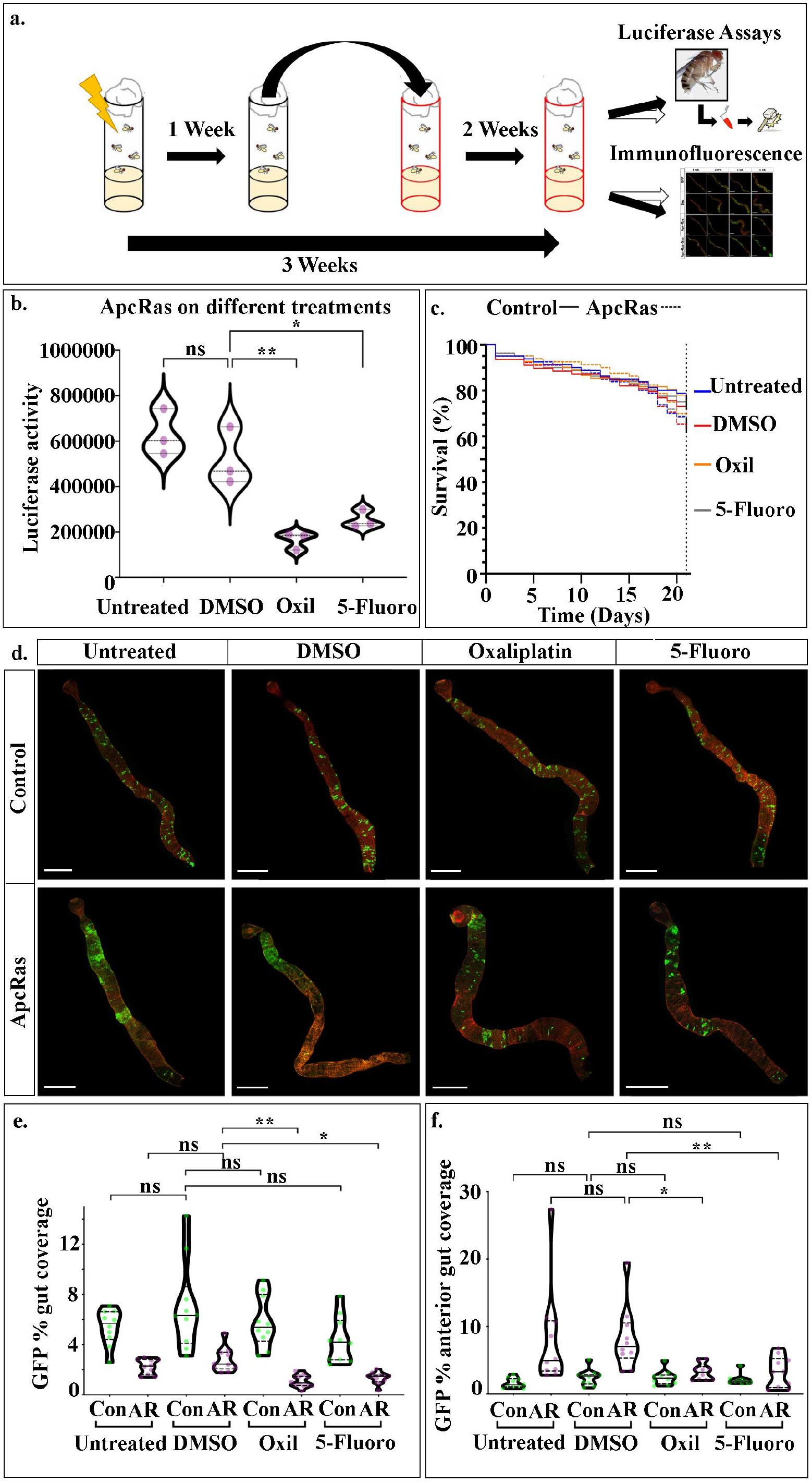
Treatment with Oxaliplatin or 5-Fluoro decreases primary tumour burden. (**a**) Schematic of the drug treatment regimen. Flies are heat-shocked (day ‘0’) and left on untreated food for 1 week, followed by transfer to a food containing drugs for 2 weeks. The flies are then sacrificed and assayed for tumour burden. (**b**) Luciferase assays performed on 3 batches of 10 ApcRas flies 3 weeks after clone induction, treated with food either without anything added (Untreated), with DMSO alone (DMSO) or with drugs added that were dissolved in DMSO (Oxaliplatin) or (5-Fluoro). (**c)** Survival curves for control (solid lines) and ApcRas (dotted lines) flies fed on untreated food (blue) or on food with DMSO alone (red), Oxil (orange) or 5-Fluoro (grey) added with DMSO. The survival of the flies was monitored over the 21 days of each experiment. Numbers were the same for each genotype: untreated n=80, DMSO n=78, Oxil n=80, 5-Fluoro n=82. (**d**) Dissected midguts from either control or ApcRas flies stained for GFP (green) and phalloidin (red). (**e**,**f)** Quantifications of the GFP% coverage of the entire midgut (**e**) or of the just the anterior-portion of the midgut (**f**) of control (Con) or ApcRas (AR) clones at 3 weeks on the different drug regimens, n = 10 for each condition. Graphs show violin plots with the median (solid line) and upper and lower quartiles (dashed lines). Statistical analysis is Kruskall-Wallis test. ** P ≤ 0.01; * P ≤ 0.05; ns = non-significant. Scale bars = 500μm.

We also followed the % survival of both ApcRas and control flies over the 3 weeks period as another potential readout for the screen. However, we found only subtle affects with each genotype and treatment: DMSO slightly lowers the survival rate of control flies, and this is partially rescued with feeding on DMSO+Oxil or DMSO+5’Fluoro (Fig 4c); DMSO also lowers the survival rate of ApcRas flies, and this is also partially rescued by treatment with either Oxil or 5-Fluor, with Oxil bringing the % survival almost as high as control flies fed on DMSO (Fig 4c). These small changes in survival rates over the 3 week period fits with previous data showing that ApcRas flies display a 50% survival rate at 4 weeks after clone induction^19^, suggesting that flies would need to be monitored for longer than 3 weeks for any effects on survival to be seen. For this to be a robust readout of the screen, it would need to be performed on more flies, over an even longer time course. Taken together, this suggests that luciferase readings on whole flies presents a far more rapid assay to screen for putative CRC treatments in flies.

We next sought to characterise in more detail how flies bearing ApcRas clones respond to treatment with Oxaliplatin or 5-Fluoro, and determine if their effects are specific to the tumour-like ApcRas clones, by comparing with control clones. To do this, we studied the behaviour of both control and ApcRas clones in the midguts of flies fed with the different conditions, by dissecting out their midguts after 3 weeks and staining them for GFP and phalloidin, which labels F-actin and can be used to visualise the outline and overall organisation of the midgut. We found that the midgut morphology and size was not affected and the distribution and size of control clones were the same in all conditions (Fig. 4d). In contrast, while the midguts of ApcRas flies raised on untreated and DMSO food had large tumour-like clones in the anterior region of the midguts, the clones in flies fed with Oxaliplatin and 5-Fluoro were greatly reduced (Fig. 4d). Thus, the reduction in clone size and distribution appears to be specific to clones with tumour-like properties.

We quantified the behaviour on clones in the different conditions by dissecting, fixing and immunostaining the midguts from these flies, and then analysing the clonal area as a percentage of the anterior region of the midgut or of the whole midgut. Both analyses revealed a reduction in coverage by GFP positive clones by around 50% when the flies were treated with either Oxaliplatin or 5-Fluoro (Fig. 4d, e). The fact that no reduction was seen in the coverage by control clones (Fig. 4e, f) suggests that treatment with the standard chemotherapies used to treat human CRC specifically target the tumour-like ApcRas clones, without causing any significant changes to normal wildtype cells.

To look in more detail at the behaviour of ApcRas clones upon treatment with Oxaliplatin or 5-Fluoro, we next used our automated scripts to analyse the area of all the clones across the entire midgut in ApcRas conditions on each treatment. We found that many very large clones over 7,800 μm2 in size are found in both untreated ApcRas flies, or ApcRas flies treated with DMSO alone. Strikingly, after ApcRas flies have been treated with either Oxaliplatin or 5-Fluoro, almost no flies are found to contain clones over the 7,800 μm2 threshold, further confirming the efficacy of these treatments (Fig 5a). In support of this, we found that other tumour-like characteristics of ApcRas clones are reduced (Fig 5b-i); the maximum nuclear size within a clone was reduced from 986 μm^2^ to ∼680 μm^2^, and while cross sections of midguts showed that there was still some multilaying of cells within ApcRas clones, they never caused the complete luminal blockages seen when these tumour-like clones were left to grow untreated (Fig. 5 compare g,i with c,e). Taken together, these results suggest that while treatment with CRC therapeutics did not completely abrogate tumour growth, the tumour burden was dramatically reduced, and this was accompanied by a reduction in the appearance of clones bulging into the lumen of the gut.

**Figure 5.**
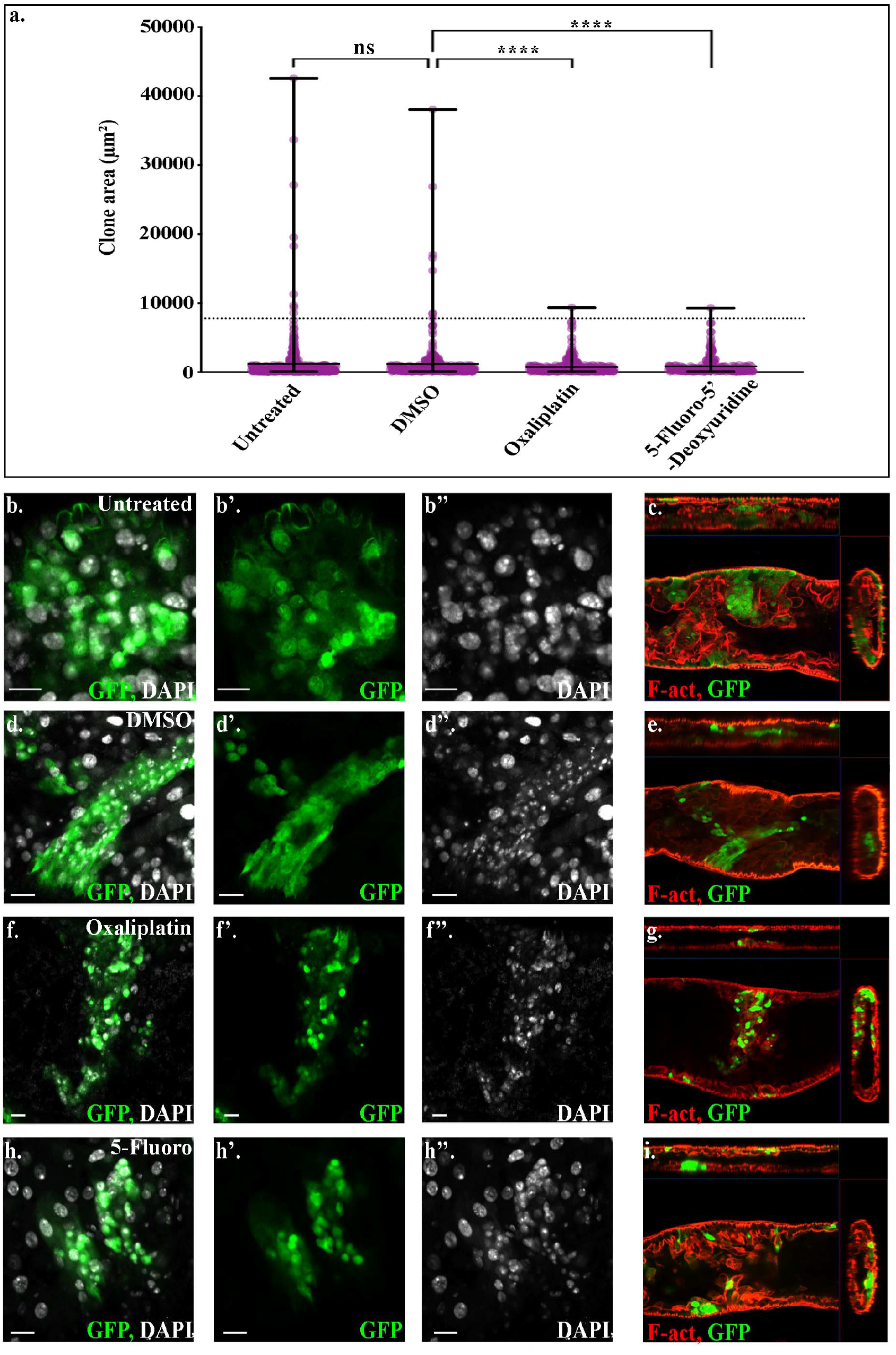
Treatment with either Oxilplatin or 5’-Fluoro results in a decrease in tumour-like clones in the ApcRas model. (**a)** Quantifications of the area of clones found in midguts of ApcRas clones on different drug regimens. Data is displayed as a scatter plot with min and max (**a**, solid horizontal lines) and a dashed line delineating 7,800 μm^2^, clones larger than which are normally only found in tumourigenic conditions. While many clones over 7,800 μm^2^ are found in ApcRas flies in untreated or DMSO conditions, when treated with Oxaliplatin or 5-Fluoro clones larger than 7,800 μm^2^ are rarely found. Statistics were performed on the number of clones found over the threshold. **b-i** Representative ApcRas clones in flies fed with untreated food (**b**,**c**) food with DMSO (**d**,**e**) with Oxilplatin (**f**,**g**) or with 5’-Fluoro (**h**,**i**). **b**,**d**,**f**,**h** are longitudinal sections and **c**,**e**,**g**,**i** orthogonal views. (**c-i**) ApcRas flies treated with Oxaliplatin or 5-Fluoro show reduced formation of tumour-like clones, multilayering and luminal blockages. Statistical analysis is Kruskall-Wallis test. **** P ≤ 0.0001; ns = non-significant. Scale bars = 50

## Discussion

To date, current drug discovery pipelines have had very limited success in delivering drugs identified *in vitro*, into the clinic for anti-cancer therapy^33,34^. With research in *Drosophila* providing the first glimpse into the mechanism of action of many human cancer-related proteins, it is now emerging as an important new platform for anti-cancer drug discovery^35^. Rapid and cost effective, its key advantage over classic *in vitro* screens, is it can identify drugs that can specifically target tumours *in vivo*. This permits assessment of drug absorption and toxicity, reducing the possibility of false positives, as well as assaying drug effects on tumour cells which are maintaining key interactions occurring during normal disease progression, such as with the tumour microenvironment. Moreover, the fast genetic manipulation of *Drosophila* presents the possibility to personalize transgenic cancer models, in which mutations are directly based on individual tumour genome profiles. Such models have been used to analyse a collection of drugs and assess the mechanisms of emerging drug resistance^36^. Remarkably, the generation of a personalized model of a treatment-resistant metastatic KRAS-mutant colorectal cancer in *Drosophila* larvae has led to the identification of a drug combination, trametinib and zoledronate, that has been used as a treatment for the patient^37^, highlighting the potential of *Drosophila* as a model for personalized medicine. Here, we have developed sensitive high-throughput assays, to rapidly and accurately assay for tumour burden in a *Drosophila* model for benign CRC, that has previously been shown to recapitulate many hallmarks of the human disease^19^. We show that it responds to treatment with standard chemotherapies used to treat human CRC, suggesting that compounds identified in future screens in this model may be transferable to humans.

Introducing two reporters into the system, firefly luciferase and GFP, makes the *Drosophila* ApcRas model amenable to two distinct methods of screening. The first, assaying for luciferase activity in whole flies, enables high-throughput batch processing of whole flies, to identify overall changes in tumour burden. The second method relies on the visualisation and quantification of the GFP reporter. It not only enables the verification of changes detected by luciferase assays, but gives a more in-depth analysis of individual clone behaviour in a semi high-throughput capacity. Excitingly, this suggests that screening batches of flies with luciferase assays could work as a primary screen, and positive candidates could be followed up in a second screen where midguts are dissected and analysed using our high-throughput scripts. Here we show that a *Drosophila* model for benign CRC rapidly responds to treatment with standard chemotherapy drugs, and that the response can reliably and reproducibly be assessed in a high-throughput manner. This process is highly scalable to the screening of thousands of flies, opening the door for drug screens to test either single compounds, or multiple compounds in combination. As *Drosophila* are highly genetically tractable, once a drug or drug combination has been identified, this could later be combined with a genetic screen to identify the drugs *in vivo* targets. Given the current need for new therapies for CRC that have less side effects and are more effective than conventional ones, the ApcRas *Drosophila* model presents an exciting new avenue, at a fraction of both cost and time of vertebrate models.

## Acknowledgements

We are thankful to the rest of the Campbell lab, the Casali lab and the Bulgakova lab for helpful discussions. We thank the Bloomington stock centre for kindly sending us reagents. This work was supported by an EPSRC DTP studentship (JA), a Wellcome Trust/Royal Society Sir Henry Dale Fellowship (KC, Grant number 204615/Z/16/Z) and a grant from the Spanish Ministry of Science and Innovation (AC, PID2019-1946GB-100).

## References

1. in World Cancer Reports. (eds. C.P. Wild, E. Weiderpass & B.W. Steward) (International Agency for Research on Cancer, Lyon, France; 2020).

2. Geng, F., Wang, Z., Yin, H., Yu, J. & Cao, B. Molecular Targeted Drugs and Treatment of Colorectal Cancer: Recent Progress and Future Perspectives. Cancer Biother Radiopharm 32, 149–160 (2017).

3. Tredan, O., Galmarini, C.M., Patel, K. & Tannock, I.F. Drug resistance and the solid tumor microenvironment. J Natl Cancer Inst 99, 1441–1454 (2007).

4. Marmol, I. et al. Alkynyl gold(I) complex triggers necroptosis via ROS generation in colorectal carcinoma cells. J Inorg Biochem 176, 123–133 (2017).

5. Rowehl, R.A. et al. Establishment of highly tumorigenic human colorectal cancer cell line (CR4) with properties of putative cancer stem cells. PLoS One 9, e99091 (2014).

6. Li, K., Zhou, Z.Y., Ji, P.P. & Luo, H.S. Knockdown of beta-catenin by siRNA influences proliferation, apoptosis and invasion of the colon cancer cell line SW480. Oncol Lett 11, 3896–3900 (2016).

7. Zhang, Y. et al. Overexpression of WNT5B promotes COLO 205 cell migration and invasion through the JNK signaling pathway. Oncol Rep 36, 23–30 (2016).

8. Konturek, P.C., Burnat, G., Rau, T., Hahn, E.G. & Konturek, S. Effect of adiponectin and ghrelin on apoptosis of Barrett adenocarcinoma cell line. Dig Dis Sci 53, 597–605 (2008).

9. Luca, A.C. et al. Impact of the 3D microenvironment on phenotype, gene expression, and EGFR inhibition of colorectal cancer cell lines. PLoS One 8, e59689 (2013).

10. Zanoni, M. et al. 3D tumor spheroid models for in vitro therapeutic screening: a systematic approach to enhance the biological relevance of data obtained. Sci Rep 6, 19103 (2016).

11. Horvath, P. et al. Screening out irrelevant cell-based models of disease. Nat Rev Drug Discov 15, 751–769 (2016).

12. Pandey, U.B. & Nichols, C.D. Human disease models in Drosophila melanogaster and the role of the fly in therapeutic drug discovery. Pharmacol Rev 63, 411–436 (2011).

13. Bangi, E. A Drosophila Based Cancer Drug Discovery Framework. Adv Exp Med Biol 1167, 237–248 (2019).

14. Markstein, M. et al. Systematic screen of chemotherapeutics in Drosophila stem cell tumors. Proc Natl Acad Sci U S A 111, 4530–4535 (2014).

15. Casali, A. & Batlle, E. Intestinal stem cells in mammals and Drosophila. Cell Stem Cell 4, 124–127 (2009).

16. Gonzalez, C. Drosophila melanogaster: a model and a tool to investigate malignancy and identify new therapeutics. Nat Rev Cancer 13, 172–183 (2013).

17. Villegas, S.N. One hundred years of Drosophila cancer research: no longer in solitude. Dis Model Mech 12 (2019).

18. Jackstadt, R. & Sansom, O.J. Mouse models of intestinal cancer. J Pathol 238, 141–151 (2016).

19. Martorell, O. et al. Conserved mechanisms of tumorigenesis in the Drosophila adult midgut. PLoS One 9, e88413 (2014).

20. Wang, C. et al. APC loss-induced intestinal tumorigenesis in Drosophila: Roles of Ras in Wnt signaling activation and tumor progression. Dev Biol 378, 122–140 (2013).

21. Suijkerbuijk, S.J., Kolahgar, G., Kucinski, I. & Piddini, E. Cell Competition Drives the Growth of Intestinal Adenomas in Drosophila. Curr Biol 26, 428–438 (2016).

22. Ngo, S., Liang, J., Su, Y.H. & O’Brien, L.E. Disruption of EGF Feedback by Intestinal Tumors and Neighboring Cells in Drosophila. Curr Biol 30, 1537–1546 e1533 (2020).

23. Dar, A.C., Das, T.K., Shokat, K.M. & Cagan, R.L. Chemical genetic discovery of targets and anti-targets for cancer polypharmacology. Nature 486, 80–84 (2012).

24. Lee, T. & Luo, L. Mosaic analysis with a repressible cell marker for studies of gene function in neuronal morphogenesis. Neuron 22, 451–461 (1999).

25. Markstein, M., Pitsouli, C., Villalta, C., Celniker, S.E. & Perrimon, N. Exploiting position effects and the gypsy retrovirus insulator to engineer precisely expressed transgenes. Nat Genet 40, 476–483 (2008).

26. Campbell, K. et al. Collective cell migration and metastases induced by an epithelial-to-mesenchymal transition in Drosophila intestinal tumors. Nat Commun 10, 2311 (2019).

27. Adams, J., Casali, A. & Campbell, K. Methods to generate and assay for distinct stages of cancer metastasis in adult Drosophila melanogaster. Methods Mol Biol (2020).

28. Hudry, B., Khadayate, S. & Miguel-Aliaga, I. The sexual identity of adult intestinal stem cells controls organ size and plasticity. Nature 530, 344–348 (2016).

29. Zipper, L., Jassmann, D., Burgmer, S., Gorlich, B. & Reiff, T. Ecdysone steroid hormone remote controls intestinal stem cell fate decisions via the PPARgamma-homolog Eip75B in Drosophila. Elife 9 (2020).

30. Kuipers, E.J. et al. Colorectal cancer. Nat Rev Dis Primers 1, 15065 (2015).

31. Arango, D. et al. Molecular mechanisms of action and prediction of response to oxaliplatin in colorectal cancer cells. Br J Cancer 91, 1931–1946 (2004).

32. Wilson, P.M. et al. Novel opportunities for thymidylate metabolism as a therapeutic target. Mol Cancer Ther 7, 3029–3037 (2008).

33. Munos, B. Lessons from 60 years of pharmaceutical innovation. Nat Rev Drug Discov 8, 959–968 (2009).

34. Ocana, A., Pandiella, A., Siu, L.L. & Tannock, I.F. Preclinical development of molecular-targeted agents for cancer. Nat Rev Clin Oncol 8, 200–209 (2010).

35. Yadav, A.K., Srikrishna, S. & Gupta, S.C. Cancer Drug Development Using Drosophila as an in vivo Tool: From Bedside to Bench and Back. Trends Pharmacol Sci 37, 789–806 (2016).

36. Bangi, E., Murgia, C., Teague, A.G., Sansom, O.J. & Cagan, R.L. Functional exploration of colorectal cancer genomes using Drosophila. Nat Commun 7, 13615 (2016).

37. Bangi, E. et al. A personalized platform identifies trametinib plus zoledronate for a patient with KRAS-mutant metastatic colorectal cancer. Sci Adv 5, eaav6528 (2019).

